# Interspecific Hybrids Show a Reduced Adaptive Potential Under DNA Damaging Conditions

**DOI:** 10.1101/2020.03.01.971416

**Authors:** Carla Bautista, Souhir Marsit, Christian R Landry

## Abstract

Hybridization may increase the probability of adaptation to extreme stresses. This advantage could be caused by an increased genome plasticity in hybrids, which could accelerate the search for adaptive mutations. High ultraviolet (UV) radiation is a particular challenge in terms of adaptation because it affects the viability of organisms by directly damaging DNA, while also challenging future generations by increasing mutation rate. Here we test if hybridization accelerates adaptive evolution in response to DNA damage, using yeast as a model. We exposed 180 populations of hybrids between species (*Saccharomyces cerevisiae* and *Saccharomyces paradoxus*) and their parental strains to UV mimetic and control conditions for approximately 100 generations. Although we found that adaptation occurs in both hybrids and parents, hybrids achieved a lower rate of adaptation, contrary to our expectations. Adaptation to DNA damage conditions comes with a large and similar cost for parents and hybrids, suggesting that this cost is not responsible for the lower adaptability of hybrids. We suggest that the lower adaptive potential of hybrids in this condition may result from the interaction between DNA damage and the inherent genetic instability of hybrids.

## Introduction

Heterogeneous environments constantly challenge organisms by changing which phenotypes are optimal. Understanding which mechanisms accelerate or slow down adaptation to environmental heterogeneity is a central question in evolutionary biology (Bleuven & Landry, 2016). If changes are rapid and drastic, populations may collapse before adapting, unless populations move to another environment. These movements often impact geographical distributions, which can promote encounters between incipient species, enhancing the probability of hybridization and introgression (Gómez et al., 2015).

Although hybridization can have many negative impacts on the long-term genetic integrity of some taxa (Lowe et al., 2015), it can also create favorable conditions for rapid adaptive evolution. The adaptive role of hybridization has been shown in the colonization of new niches (Gallego-Tévar et al., 2018; Lewontin & Birch, 1966), during speciation (Schumer et al., 2014), and during adaptive radiation (Ballerini, 2012). Prime examples include Darwin finches, which benefited from hybridization during adaptation to the adverse climatic conditions caused by the exceptionally severe El Niño event (Grant & Grant, 1996). Another example is increased invasiveness of plant species (Schierenbeck & Ellstrand, 2009). For instance, rise in sea level is associated with recurrent hybridization between native species and invasive ones of the *Spartina* genus in coastal marshes (Gallego-Tévar et al., 2018, 2019). In such cases, hybridization may facilitate adaptation through genomic admixture and its associated increase in phenotypic diversity, which could be particularly critical in extreme environments (Grant & Grant, 1996; Heil et al., 2017; Lexer et al., 2003; Martin et al., 2006). In addition, hybrids often show phenotypes outside of the range observed in parental species because of transgressive segregation or heterosis, and these phenotypes may be adaptive (Landry et al., 2007; Nolte & Sheets, 2005; Rieseberg et al., 1999; Vega & Frey, 1980).

Another important feature that may influence the evolution of hybrids is their inherent genomic instability, which can accelerate adaptation (Taddei et al., 1997). Genomic instability includes higher rate of DNA damage (Herbst et al., 2017), chromosomal rearrangements (Baack & Rieseberg, 2007), gene and chromosome copy number variation (Dion-Côté & Barbash, 2017) and the multiplication of transposable elements (Guerreiro, 2014). Instability often leads to aneuploidy, which has also been shown to be a mechanism of adaptation in stressful conditions (Sunshine et al., 2015), and hybrids may be particularly prone to producing aneuploid progeny (Gilchrist & Stelkens, 2019). Another consequence of genomic instability is changes in ploidy. Hybridization followed by whole genome duplication giving rise to polyploid hybrids has been observed in plants, animals and fungi (Alves et al., 2001; Marcet-Houben & Gabaldón, 2015; Soltis & Soltis, 2009). Because polyploidy itself could increase the rate at which beneficial mutations are acquired (Selmecki et al., 2015), hybrids that polyploidize could have access to this feature as well. Indeed, hybridization followed by polyploidization has been directly related with adaptive diversification in plants (Alix et al., 2017) and in fish (Saitoh et al., 2010).

There are likely limits to the adaptive potential of hybrids caused by their enhanced genomic instability. For instance, genomic changes of large effects, such as changes in ploidy, may lead to hybrid inviability (Burton & Husband, 2000). Similarly, the increase in mutation rate in hybrids (Xie et al., 2016) could lead to an increased acquisition rate of deleterious mutations. Indeed, *Escherichia coli* strains with a 100-fold increase in the mutation rate experience a reduction in adaptation in many environments (Sprouffske et al., 2018). In addition, the type of mutations that contribute to adaptation in hybrids may be particularly deleterious in other environments, leading to strong trade-offs. This is the case for aneuploid yeast strains, which typically show a large condition-specific response due to pleiotropic effects (Sunshine et al., 2015). Finally, the accelerated rate of evolution could lead to the rapid accumulation of conditionally neutral mutations (Cooper, 2014), leading to fitness trade-offs in other environments and thus limiting the long-term potential of hybrids.

*Saccharomyces* yeast species have been widely used in experimental research due to their dual sexual and asexual mode of reproduction, their short doubling time and their deep genetic characterization (Replansky et al., 2008). They have been used to study adaptation to high salinity (Dhar et al., 2011), to extreme temperatures (Salvadó et al., 2011), to dehydration (Khroustalyova et al., 2019) and to heavy metals (Adamo et al., 2012). Despite the presence of interspecific reproductive barriers, introgression among yeast species has been observed in nature (Barbosa et al., 2016; Leducq et al., 2016) and in industrial conditions (Lopandic, 2018). Evidence for the superiority of hybrids in response to extreme stress also comes from experimental evolution. For instance, Stelkens *et al*. (2014) showed that yeast hybrids are more likely to be evolutionarily rescued than parental species in stressful environments.

A prominent environmental stressor is UV radiation, which has been related with some extinctions as in the Devonian-Carboniferous boundary period (Marshall et al., 2020). Nowadays, it has been intensified with human-induced environmental change due to, among other factors, the depletion of the ozone layer (Caldwell & Flint, 1994). Radiation represents a particular challenge in terms of adaptability by affecting DNA integrity and by increasing mutation rates (Felkner & Kadlubar, 1968). Here we examine the rate of adaptation of two yeast species, *Saccharomyces cerevisiae* and *Saccharomyces paradoxus*, and of their hybrids, in the face of an environmental stressor, a DNA damaging agent mimicking the effects of UV radiation. These two species diverge by about 15% at the nucleotide level (Cliften et al., 2001) and naturally hybridize in the wild (Barbosa et al., 2016), although the F1 diploid hybrid is almost completely sterile (Murphy et al., 2006). We hypothesize that, because of the evolutionary potential of hybridization brought by genomic instability, hybrids will adapt faster than parental species. On the other hand, DNA damage may enhance genetic instability to a point where it prevents adaptation of the hybrid populations. To test these hypotheses, we experimentally evolved 180 populations of diploid hybrids and diploid parental strains through approximately 100 generations in UV-mimicking conditions and matching control conditions.

## Materials and Methods

### Yeast Strains and Media

The *S. cerevisiae* and *S. paradoxus* strains used in this study derive from natural strains LL13_054 and MSH-604 that were isolated in natural forests in North America. The *HO* locus was replaced by homologous recombination with different resistance cassettes to prevent mating type switching in haploids (Hygromycin B and Nourseothricin) as described in Güldener et al. (1996). The haploid yeast strains used (described in Table S1) were LL13_054 MATa HO:: Hygromycin B and LL13_054 MATα HO::Nourseothricin for *S. cerevisiae* and MSH-604 MATα HO::Nourseothricin and MSH-604 MATa HO::Hygromycin B for *S. paradoxus* (Charron et al., 2019; Leducq et al., 2016). As a neutral growth condition, cells were grown in YPD (1% yeast extract, Fisher BioReagents™, USA; 2% tryptone, BioShop^®^, Canada; and 2% D-glucose, BioShop^®^, Canada). The change in the batch of the yeast extract (Fisher BioReagents™, USA) used during the experimental evolution may be responsible for the slight decrease in the growth rate we observed in control conditions during this evolution experiment.

### Experimental Crosses

All incubation steps were performed at 25 °C. Haploid strains were grown overnight in 5 mL of YPD. Pre-cultures were diluted to OD_595_ of 1.0 in 500 μL aliquots. The aliquots from pairs of strains to be crossed were mixed in a tube and 5 μL were used to inoculate 200 μL of YPD medium in 30 replicates. Therefore, all starting diploid populations derive from independent mating events. Mixed haploid strains were incubated for 6 h after which 5 μL of the mating cultures were spotted on a double selection YPD solid medium (100 μg/mL of Nourseothricin and 250 μg/mL of Hygromycin B). From each of the 30 spots per genotype, 1 colony was picked as a founder population for the evolution experiment, resulting in 30 independent lines for each of the 3 genotypes (parental species 1: *S. cerevisiae*, parental species 2: *S. paradoxus*, and their hybrids).

### UV Mimetic Tolerance Assays

In order to reproduce UV radiation conditions, we used 4-Nitroquinoline 1-oxide (4-NQO) (Sigma–Aldrich, cat. no. N8141, batch #WXBC3635V), a UV mimetic molecule (Felkner & Kadlubar, 1968). 4-NQO targets DNA by forming bulky adducts, which are formed by covalently attached bases, disrupting base-pairing and altering DNA structure (Felkner & Kadlubar, 1968). 4-NQO was dissolved in dimethyl sulfoxide (DMSO) at a concentration of 400 mM in individual aliquots that were stored at −20°C. To estimate cell tolerance before experimental evolution, 20 random lines of our three genotypes were inoculated in liquid cultures with gradually increasing 4-NQO concentrations (YPD + 4 μM, 8 μM, 16 μM, 24 μM, 32 μM and 40 μM of 4-NQO) and growth was monitored for 24 h.

### Experimental Evolution

30 parallel lines for each genotype were pre-cultured in 1 mL of YPD liquid cultures in 96 deep-well plates (2 mL) and incubated for 24 h at 25 °C. Subsequently, 20 μL of these 90 parallel lines (Fig. S1) were transferred in 96-well flat-bottomed culture plates with 180 μL of media (YPD or YPD + 4-NQO), resulting in an initial OD_595_ of approximately 0.1. The borders of the plates did not contain strains and were filled with sterile media to avoid border effects related to evaporation. Overall, a total of 180 cultures were maintained in parallel, 30 replicates for each of the 3 genotypes in YPD and 30 replicates for each of the 3 genotypes in YPD + 4-NQO (Fig. S1). Every 24 h, each culture was diluted approximately 30-fold by transferring 6 μL of grown culture into 194 μL of fresh culture medium to initiate a new round of growth at an OD_595_ starting at about 0.03. A total of 21 such transfer cycles were carried out, resulting in approximately 100 generations (each transfer cycle involved approximately ~ 5 generations in rich conditions, Fig. S2). Incubation at 25 °C was performed directly in 3 temperature-controlled spectrophotometers (Infinite^®^ 200 PRO, Tecan, Reading, UK) that read the OD_595_ at intervals of 15 min. Archives were maintained for every transfer cycle by mixing 80 μL of the evolved lines with 170 μL of 80% glycerol in 96 well plates and stored in a −80°C freezer. We considered the initial growth as the growth estimated from the third cycle (T15) because we did not monitor growth continuously in the first and second cycles due to technical problems. Also, not including the first two cycles may be required because cells sensitivity to the UV mimetic chemical increased during that period (Fig. S3AB), most likely because it takes time before cells are sensitized to the stress condition.

### Fitness Assays at the End of the Experimental Evolution

Ancestor strains (n = 90) as well as the lines evolved in YPD (n = 90) and in YPD + 4-NQO (n = 90) were thawed from glycerol stocks on solid YPD omnitray plates (25 °C, 72 h). They were pre-cultured in 1 mL of YPD liquid cultures in 96 deep-well plates (2 ml) and incubated for 24 h at 25 °C. Subsequently, 20 μL of these pre-cultures were grown in 96-well flat-bottomed culture plates in 180 μL of media (YPD or YPD + 4 μM, 8 μM, 16 μM, 24 μM, 32 μM and 40 μM of 4-NQO), resulting in an initial OD_595_ of approximately 0.1. Incubation at 25 °C was performed directly in 3 temperature-controlled spectrophotometers (Infinite^®^ 200 PRO, Tecan, Reading, UK) that read the OD_595_ at intervals of 15 min.

### Statistical Analysis

Growth of each experimental line was measured as the maximum growth rate (r), the maximum slope of the growth curve fitted using the “Growthcurver” package (Sprouffske & Wagner, 2016) in R. Similarly, we calculated the carrying capacity (K), which represents the maximum population size a particular environment can support, for further correlation analysis to test its relationship with growth rate (r). We quantified the rate of adaptation as the increase of growth rate through time with both linear and non-linear models (“drc” package). We used analysis of variance (ANOVA) to test for differences in growth rates (with normal distribution fit) between groups. Paired t-tests were also used for paired data when comparing the same lines at different times (generation 0 and generation 100) or the same lines evolved in different media (control or UV mimetic). Multiple Tukey post-hoc pairwise comparisons were used to test for differences between groups. For hypothesis testing, we always considered as significant a P-value <0.05. The data was analyzed using R version 3.4.1. The code is available at <u>GitHub</u>.

## Results

### Experimental Evolution

We tested whether yeast hybrids would adapt faster than parental species to DNA stress conditions. We carried out an experiment with 90 independent populations of two parental species (*S. cerevisiae* and *S. paradoxus*) and their F1 diploid hybrid. These populations were grown for approximately 100 generations in rich media (YPD) combined with a UV radiation mimetic chemical, 4-nitroquinoline-1-oxide (4NQO), and in parallel in rich media (YPD) only as control (Fig. 1A). We refer to the *S. cerevisiae, S. paradoxus* and hybrid crosses as the three genotypes.

**Figure 1.**
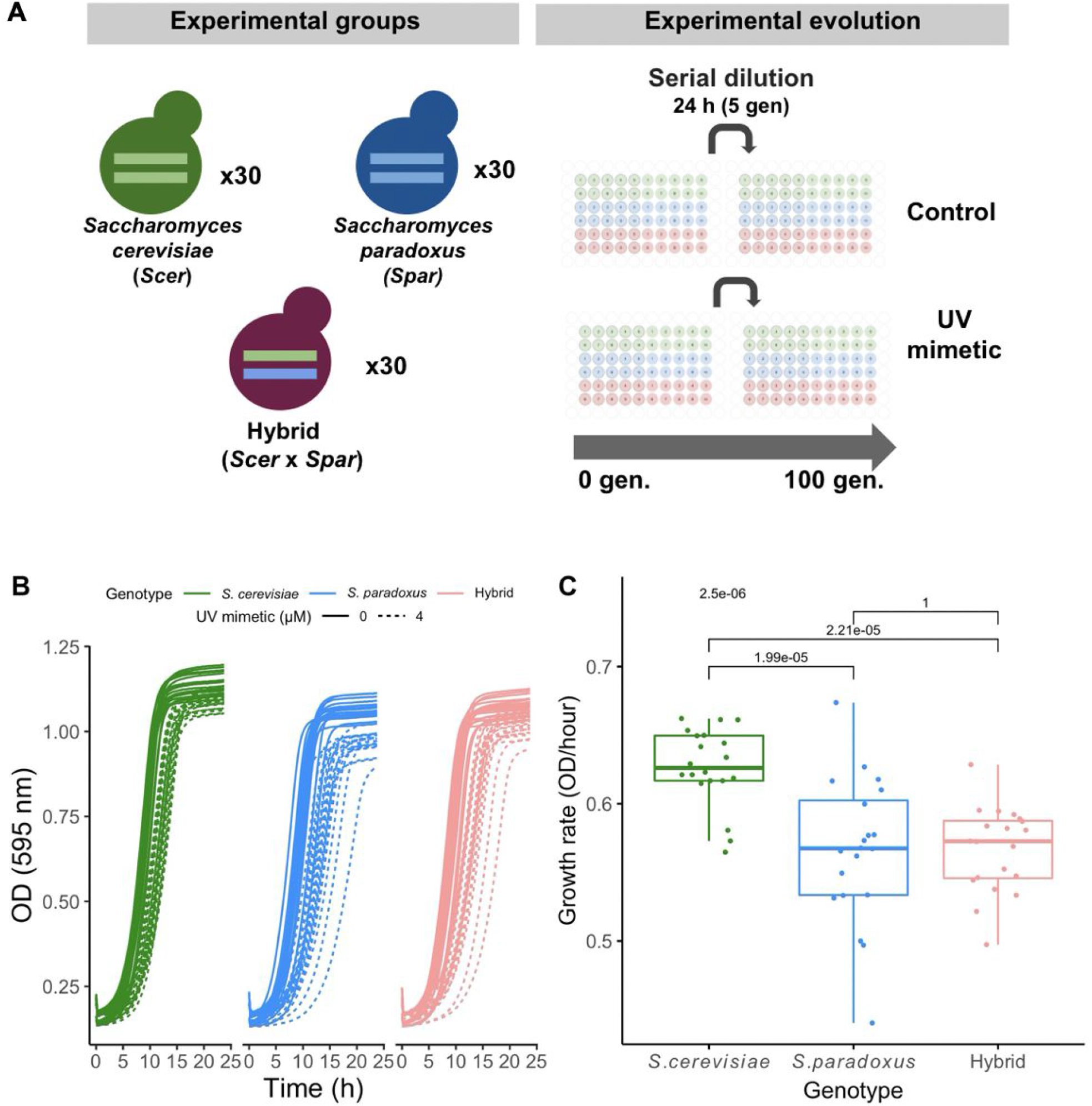
Experimental design. A) 30 independent populations of each genotype (*S. cerevisiae, S. paradoxus* and their hybrid) were evolved for approximately 100 generations in control YPD and in YPD + a UV mimetic chemical (4 μM of 4-NQO). Each 24 h, a new population was founded by transferring about 3% of the previous population to fresh media. B) Test for growth of the 3 genotypes in 4-NQO. Optical density as a function of time for 20 of the initial populations of each genotype in control conditions and in UV mimetic conditions (4 μM of 4-NQO). C) Growth rate of strains in 4 μM of 4-NQO (n = 20 populations for each genotype). P-value for ANOVA test (above) and Tukey post-hoc pairwise p-values are shown.

In order to determine the proper UV mimetic chemical concentration to use, we first performed dose–response experiments across concentrations ranging from 0 to 40 μM of 4-NQO in 20 randomly chosen populations. Growth was inhibited with increasing concentrations (Fig. S4A). We observed a steady reduction of growth rates from about 20% at 4 μM to nearly 80% at 40 μM across the three genotypes (Fig. S4B). We first tested 16 μM as concentration for experimental evolution but populations failed to grow in the second round of serial dilution (Fig. S3B). We selected 4 μM as concentration, which led to approximately 25-30% growth rate diminution for the three genotypes (Fig. 1B). In general, *S. cerevisiae* was slightly less sensitive than *S. paradoxus* and the hybrid (Fig. 1C, Fig. S4B). *S. paradoxus* showed more heterogeneity among replicates, with some of them being inferior to those of the hybrid (Fig. 1C).

By comparing the maximum growth rate in UV mimetic condition through time, we found that all 3 genotypes experienced a significant increase in growth rate (Fig. 2A) and have thus adapted. The UV mimetic adapted lines reached on average 76%. 64% and 41% of the initial growth rate in control conditions for *S. cerevisiae, S. paradoxus* and the hybrids respectively (percentages calculated using T100 in Fig. 2B). We found a slight decrease in growth rate in control conditions when comparing data gathered during experimental evolution (Fig. S5A). However, we find that this was caused by a slight change in experimental conditions during the experiment (see material and methods section) since these same strains grown in YPD once the experimental evolution ended show no significant differences (Fig. S5B). This change in media was previously remarked by Charron *et al*. (2019) in our laboratory. We measured the growth rate of the evolved strains from T100 across the gradient of UV mimetic concentrations tested above and found that fitness of evolved populations increased compared to the ancestral ones (Fig. S6). This result shows that there was a general decrease in sensitivity to the UV mimetic chemical and thus that adaptation is not specific to the concentration used for experimental evolution but extends to other concentrations as well.

**Figure 2.**
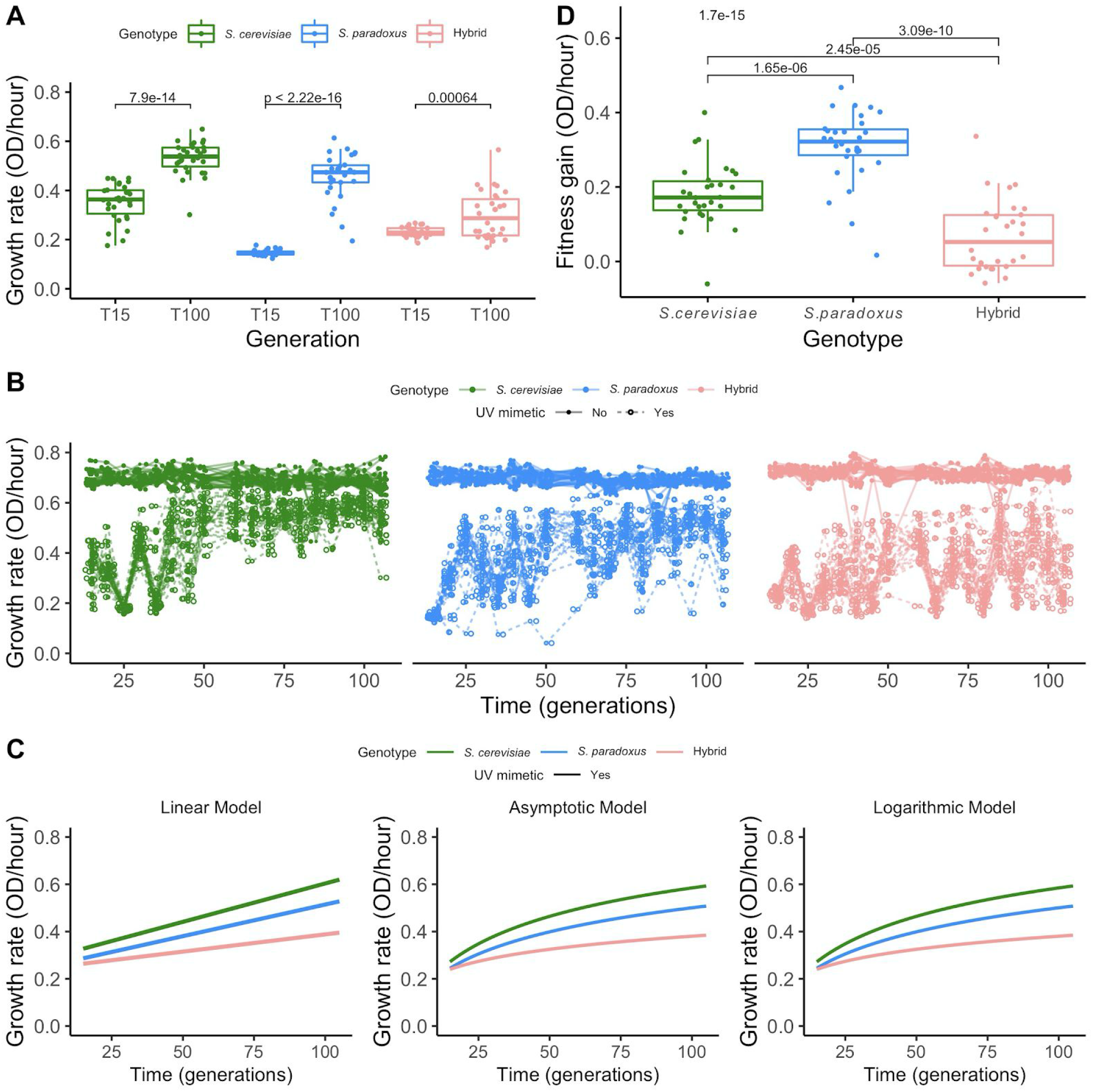
Adaptation to UV mimetic conditions. A) Growth in UV mimetic conditions for the 30 lines of the three genotypes at the initial (T15) and final (T100) time points. Paired t-tests were performed between growth rate at T15 and T100 for each genotype. P-values are shown above. B) Growth rates as a function of the number of generations. Each data point is a growth rate estimated per population per time cycle. Lines represent the evolutionary trajectories of individual replicates (n = 30 populations for each genotype). C) Statistical modeling of the growth rate of the different genotypes as a function of the number of generations is shown. Left panel corresponds to the Linear Model (n = 30 populations for each genotype), center, to the Asymptotic Model (n = 30 populations for each genotype) and right, to the Logarithmic Model (n = 30 populations for each genotype). See Table S2, S3, S4 and S5 for further details. D) Fitness gain: change in the growth rate between initial and final time points calculated by subtracting growth rate in UV mimetic conditions at T15 from the growth rate at T100 (n = 30 populations for each genotype). P-value for ANOVA test (above) and Tukey post-hoc pairwise p-values are shown.

Growth rate did not systematically increase in UV mimetic conditions. We saw paths of increased and decreased growth rate through time (i.e. a up and down pattern) (Fig. 2B). This pattern fades rapidly in *S. cerevisiae* (in about 50 generations), and then in *S. paradoxus* (in about 80 generations), with some exceptions. The hybrid experiences this pattern throughout the experiment. We fitted different models to quantify the average increase of growth rate through time (Fig. 2C, Table S2), but did not attempt to explain these oscillatory patterns with the model. We first fitted a linear model (Fig. 2C left) to model the increase in growth rate through time. Because previous studies have also shown that adaptation could be rapid at first and slow down with time, we also considered two non-linear models in which the rate decreases with time: asymptotic (Fig. 2C center) and logarithmic (Fig. 2C right) (Table S2). The comparison of Akaike Information Criterion (AIC) scores reveals a slightly better fit of the asymptotic model, followed by the logarithmic and then the linear model (Table S2). This suggests that indeed, rates of adaptation are initially high and decrease with time, as observed in other studies (de Visser & Lenski, 2002). The fit with the asymptotic model suggests also that there may be an upper limit of adaptation that would correspond to growth under control conditions, which did not improve during the experiment.

The comparison of fitness gains between the final and initial time points (Fig. 2D) shows that the extent of adaptation in the hybrid was lower than in the two parental genotypes (Anova and Tukey post-hoc pairwise comparisons, *p* = 2.45e-05 and *p* = 3.09e-10, respectively for *S. cerevisiae* and *S. paradoxus*), which rejects our hypothesis that hybrids would show a greater rate of adaptation. We noticed the same result when observing the slopes that reflect the rate of adaptation through time. The hybrid shows a slope in UV mimetic conditions that is significantly shallower than both parental species in the linear model (Fig. 2C left, Table S3: *p* < 1.26e-14 and *p* < 1.31e-07, respectively for *S. cerevisiae* and *S. paradoxus*). Moreover, the increase in the growth rate is significantly lower for the hybrid also in the logarithmic model (Fig. 2C right, Table S4: *p* = 6.516e-16 and *p* = 3.968e-09, respectively for *S. cerevisiae* and *S. paradoxus*). However, in the asymptotic model, the increase in growth rate is only significantly different when comparing the hybrid and *S. paradoxus* (Fig. 2C center, Table S5: *p* = 0.03229). Nevertheless, this model also reveals that the maximum growth rate achieved was significantly lower for the hybrid than for both parents (Fig. 2C center, Table S5: *p* = 7.558e-15 and *p* = 4.307e-05, respectively for *S. cerevisiae* and *S. paradoxus*). All models therefore support a lower rate of adaptation in hybrids. This result is also seen when we exposed the 3 genotypes to a gradient of concentration of the UV mimetic chemical in which the increased tolerance is less pronounced in the hybrid at high doses (Fig. S6).

Finally, we analyzed the correlation between the two parameters that characterize density-dependent population growth: the maximum growth rate per individual (r), as used above, and the carrying capacity (K), which is the optical density in saturated cultures. It was recently shown that the *r–K* correlation is negative in optimal conditions and positive in stressful conditions (Wei & Zhang, 2019). Accordingly, we hypothesized that the r-K correlation would be positive in UV mimetic conditions, and that its value would decrease as the strains adapted because the extent of stress would decrease. We found that the values of the *r*–*K* correlations are negative in control conditions and positive in UV mimetic conditions (Fig. S7). The correlation in UV mimetic conditions increases slightly in the hybrid while it remains stable (*S. cerevisiae*) or decreases (*S. paradoxus*) in parental species (Fig. S7). This supports that adaptation was less pronounced in the hybrid compared to the parents, although the differences are subtle.

### Cost of Adaptation

We tested whether adaptation to UV mimetic conditions would lead to a fitness trade-off in other conditions, and if this trade-off was stronger for hybrids compared to parents. Such a trade-off could come from the cost of adapting to UV mimetic conditions, or to the accumulation of mutations that are neutral in the stress conditions but would be deleterious in the control conditions, both of which would be visible in control conditions. To test this, we measured the growth rate of all the strains in control conditions (YPD). As shown above, we observed no growth rate improvement in control conditions for the strains evolved in control conditions compared to the ancestors grown in the same conditions (Fig. 3A) (Paired t-tests, *p* = 0.68, *p* = 0.66, *p* = 0.089, respectively for *S. cerevisiae, S. paradoxus* and hybrid). However, strains evolved in UV mimetic conditions grew slower than both ancestors and strains evolved in control conditions (Fig. 3A) when grown in control conditions. Therefore, all 3 genotypes showed this fitness trade-off once adapted to UV mimetic conditions. Hybrid genotypes only showed a slightly stronger trade-off than *S. paradoxus* but not compared to *S. cerevisiae* (Fig. 3B) (Anova and Tukey post-hoc pairwise comparisons, *p* = 0.0216 and *p* = 0.947, respectively). The proposed linear model which refers to the extent of the trade-off as a function of the extent of adaptation (Fig. 3C) explained only 9% of the variability observed but it is still significant (p = 0.02). However, we observed a correlation between the extent of the trade-off and the extent of adaptation (Fig. S8: Spearman’s rank correlation coefficient r = 0.33, *p* = 0.0016). There were no significant differences among genotypes in the trade-off/adaptation ratio, supporting the observation that adaptation in hybrids is not more costly than in the parents (Fig. 3D). Some strains showed negative trade-off/adaptation ratios (Fig. 3D) and those come either from negative trade-off or null or slight loss of fitness (no adaptation, values below 0). This is caused by the fact that 12.22% of the strains evolved in the UV mimetic conditions have lower growth rates at the end of the experiment than their ancestors. Almost all of these (10/11) values belong to hybrids, consistent with their lower rate of adaptation.

**Figure 3.**
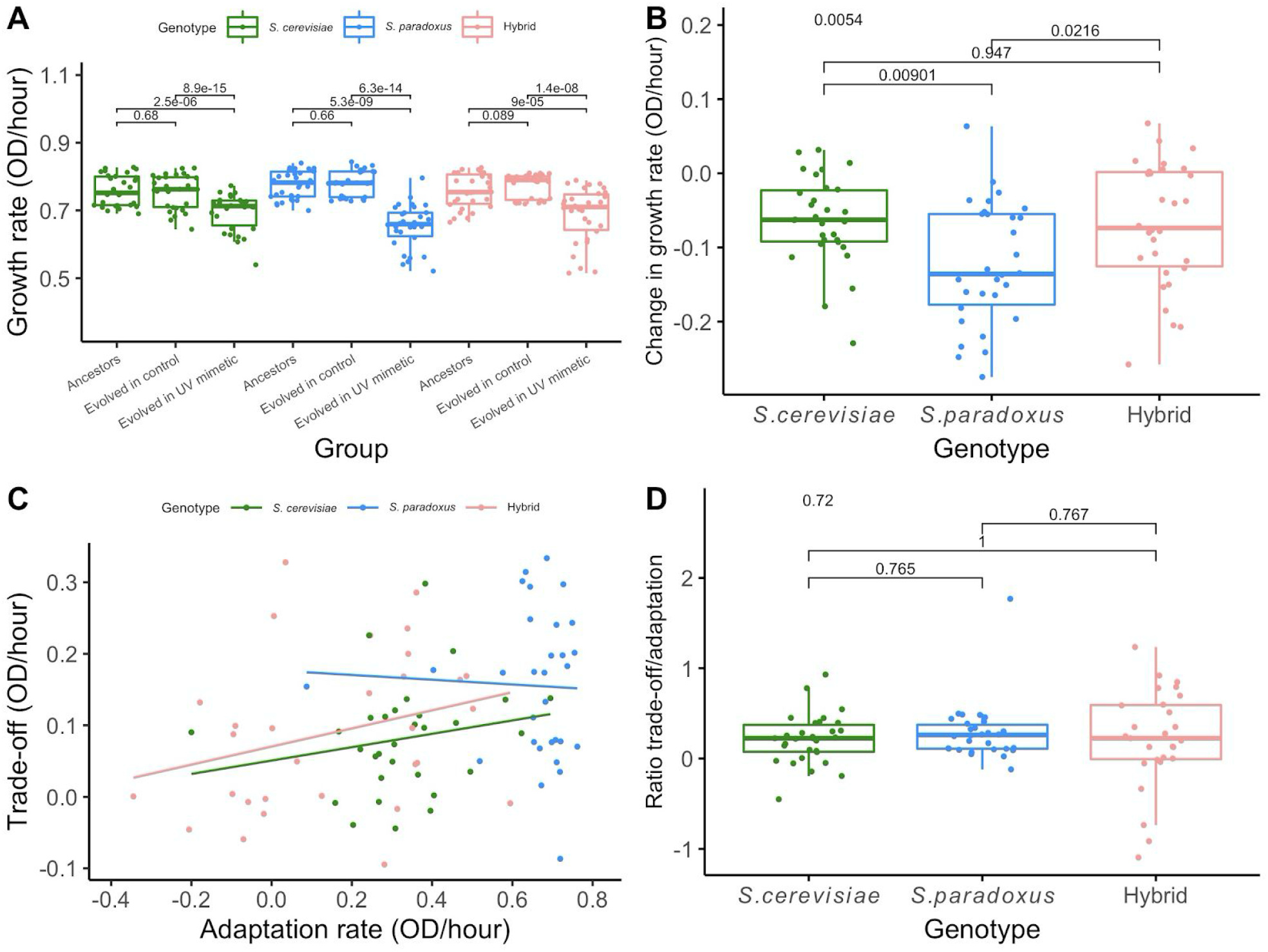
Strains adapted to a UV mimetic chemical show a trade-off under control conditions. A) Growth rate in control conditions of ancestral strains, of strains evolved in controlled conditions (100 generations) and of strains evolved in UV mimetic conditions (100 generations). Paired t-tests were performed by matching individual strains (n = 30 populations for each genotype). P-values are shown above. B) Trade-off represented as a change in growth rate between strains evolved in UV mimetic conditions and their ancestors calculated by subtracting growth rate in control of ancestors from the growth rate in control of the strains evolved in UV mimetic conditions (T100). ANOVA and Tukey post-hoc pairwise comparisons were performed among genotypes (n = 30 populations for each genotype). P-value for ANOVA test (above) and Tukey post-hoc pairwise p-values are shown. C) Trade-off depends on the extent of adaptation to UV mimetic conditions. Adaptation rates, expressed as fitness increase over the experiment, were obtained by subtracting the growth in UV mimetic conditions at T15 from the growth in UV mimetic conditions at T100. Trade-off estimates were obtained by subtracting the growth rate of the strains evolved in UV mimetic conditions (T100) grown in control conditions from the growth of the ancestors grown in control conditions. Linear model of the extent of the trade-off as a function of the extent of adaptation for each genotype (n = 30 populations for each genotype). D) Cost of adaptation as the trade-off/adaptation ratio calculated by dividing the values of trade-off by the values of adaptation for each genotype. ANOVA and Tukey post-hoc pairwise comparisons were performed among genotypes (n = 30 populations for each genotype). P-value for ANOVA test (above) and Tukey post-hoc pairwise p-values are shown.

## Discussion

Identifying which factors favor adaptation to environmental stressors is an important goal in ecology and evolution. Here we tested if hybrids would adapt faster than parental species when exposed to intense stress, using yeast and UV radiation mimetic conditions as models. Our hypothesis was based on previous observations suggesting that hybrids may have an adaptive advantage when faced with stressful conditions (Lopandic, 2018; Stelkens et al., 2014). Previous studies reported rapid adaptation in yeast (50-300 generations) in stresses such as salt, copper and ethanol (Adamo et al., 2012; Dhar et al., 2011; Khroustalyova et al., 2019; Salvadó et al., 2011) and also to UV radiation both in natural strains (Lidzbarsky et al., 2009; Pulschen et al., 2015) and in laboratory conditions (Lawrence & Christensen, 1976). We found that all three genotypes adapted to UV mimetic conditions, but the rate of adaptation and the extent achieved in 100 generations were lower for hybrids than for parental species. Moreover, hybrid replicates were more variable in their adaptive trajectories. This greater variation might be explained by their increased instability and larger access to mutations and/or genotypes, resulting from the fact that they derive from two different genomes.

We saw that growth rates fluctuate from one round of dilution to the next at the beginning of the experiment. The cause of this pattern is unclear but one possibility is that strains are initially very sensitive to 4-NQO but this sensitivity may vary as a function of the growth phase. For instance, cells could be less sensitive as they enter stationary phase (Siede & Friedberg, 1990). The growth rate in the next cycle would therefore be dependent on the phase in which cells were in the previous cycle. This effect would temper when sensibility decreases with adaptation. We see a significant association between the growth rate and the final OD of the previous cycle (Spearman’s rank correlation coefficient r = 0.46, *p* < 1e-16). Another possibility would be that adaptive mutations would occur and disappear in the next round through dilution. However, this is very implausible because the bottlenecks still represent relatively large population sizes and this effect would unlikely occur in a synchronized fashion across replicate lines. Nevertheless, the increased mutation rate caused by the UV mimetic may interact with drift in such a way that deleterious mutations may reach high frequencies after a bottleneck. This remains to be examined.

Why do hybrids show a slower rate of adaptation than parental strains? One possibility would be that hybrids are more sensitive to the chemical 4-NQO, so it makes them incapable of adapting to this media. This is unlikely since the hybrid genotype is less sensitive than *S. paradoxus* at the concentration used. We rather hypothesize that in the specific case of stress caused by DNA damage, hybrids may be at a disadvantage because they are genetically unstable (Baack & Rieseberg, 2007; Guerreiro, 2014; Marfil et al., 2006; Morales & Dujon, 2012) and DNA damaging agents may further enhance this instability, preventing the occurrence or fixation of adaptive mutations. Consistent with the instability of hybrids, recent study of mutation accumulation in yeast reported that hybrids from more divergent parents lines were lost at a greater rate than the less divergent ones (Charron et al., 2019). As the loss occurred in the first 250 generations, the authors suggested that it was mostly due to the genomic instability that arises rapidly after hybridization rather than spontaneous mutations. Thus, we suggest that the lower adaptive potential of our hybrids in UV mimetic conditions may result from the interaction between DNA damage and the inherent genetic instability of hybrids. To support this hypothesis, it would be necessary to compare the rate of adaptation of hybrids in other DNA damaging conditions and other stresses non-related to DNA damage, and also to confirm that genome instability is enhanced in hybrids. It would also be useful to identify the mechanistic basis of adaptation to UV mimetic chemicals in hybrids and parental species, as it is possible that hybrids do not have access to the same adaptive mutations as parental species. Some mechanisms of resistance to 4-NQO have been characterized in *S. cerevisiae* and could vary with respect to hybrids, for instance through the perturbation of the proteasome (Karpov et al., 2019) and pathways involved in multi-drug resistance transporters (Rong-Mullins et al., 2018).

Adaptation to one environment can decrease fitness in another, revealing evolutionary trade-offs (Cooper & Lenski, 2000). Such trade-off can be the consequence of mutations that are beneficial in stressful environments but detrimental in non-selective environments, also called fitness cost. For instance, Arctic *Daphnia* naturally exposed to UV evolved a melanic morph that is more resistant to UV and that is competitively inferior to the non-melanic ones in laboratory conditions (Hessen, 1996). Such trade-off can limit the long-term success of populations that adapt to acute stress because they lead to specialized genotypes (Anderson et al., 2013) that would lose their advantage when normal conditions return. The accumulation of neutral mutations in strong selective environments, but that are deleterious in non-selective environments, can also cause trade-offs (Mee & Yeaman, 2019). The latter mechanism may be accelerated in the presence of UV mimetic chemicals and lead to genomic decomposition, as seen in mutator strains (Couce et al., 2017). All strains showed a significant fitness reduction in non-selective control conditions once adapted to UV mimetic conditions. We also found that across strains, the extent of trade-off is correlated with the extent of adaptation, an observation that is more consistent with an actual cost of adaptation than with the accumulation of conditionally deleterious mutations, which, in principle, should be independent from the extent of adaptation. We did not find that the hybrids suffer from a higher cost than the parental species. It therefore remains unclear whether the higher cost to adaptation and more accumulation of mutations in the hybrids contribute to limiting their rate of adaptation.

Experimental evolution comes with limitations. For instance, serial transfers produce populations that are usually subject to bottlenecks (Wahl et al., 2002). In our study, serial transfers produced an approximately 30-fold decrease in population sizes with varying severity. As a result, we observed some bottlenecks, especially among the first generations when strains are less acclimated and adapted, which made the final OD before transfer more variable due to a delay in reaching stationary phase. However, these bottlenecks did not reach critically small population sizes and are typical of experimental evolution. Otherwise, alternative evolutionary pathways could still be lost, limiting the number of different adaptive peaks visited (Poelwijk et al., 2007). Our data suggests that it did not prevent adaptation from occurring. Another limit is that we are studying F1 hybrids, which have not undergone sexual reproduction and cannot exploit the power of recombination to remove deleterious combination of alleles and favor the advantageous ones (Schumer et al., 2018). These experimental hybrids rely only on the loss of heterozygosity (Charron et al., 2019) for the removal of alleles. Sexually recombining hybrids may therefore not suffer from the same limits as F1 hybrids in the conditions we used. This will need to be examined in the future.

Hybridization between species can be a dead end or a stimulus for adaptation. The study of hybridization has been a prominent research theme in biology (Anderson, 1953; Dobzhansky & Pavlovsky, 1958; Grant & Grant, 1996; Mitchell et al., 2019; Seehausen, 2004) that may be particularly relevant to global climate changes and increase anthropic pressures on ecosystems. Indeed, understanding why hybridization may lead to advantages or disadvantages is crucial for the conservation of ecosystems (Becker et al., 2013; Chan et al., 2018; Hamilton & Miller, 2016), for crop improvement (Liu et al., 2005; Waara & Glimelius, 1995) and in industry (Bellon et al., 2015; Gibson & Liti, 2015; Lopandic, 2018). One of these changes could be changes in radiation that cause DNA damage. We therefore tested if hybridization can increase the rate of adaptation to such conditions. Our results show that studying hybrids in various conditions may be a powerful context in which to study the limits of hybridization as a mechanism that may promote adaptation in stress conditions.

## Acknowledgments

We thank Nadia Aubin-Horth, Angel Cisneros, Guillaume Charron and Marika Drouin for comments on the manuscript, Guillaume Charron for help with the strain construction, Alexandre Dubé and Isabelle Gagnon-Arsenault for help in performing some experiments, Angel Cisneros and Diana Ascencio for help in data analysis and members of the Landry Lab for helpful discussions.

---

**Figure S1.**
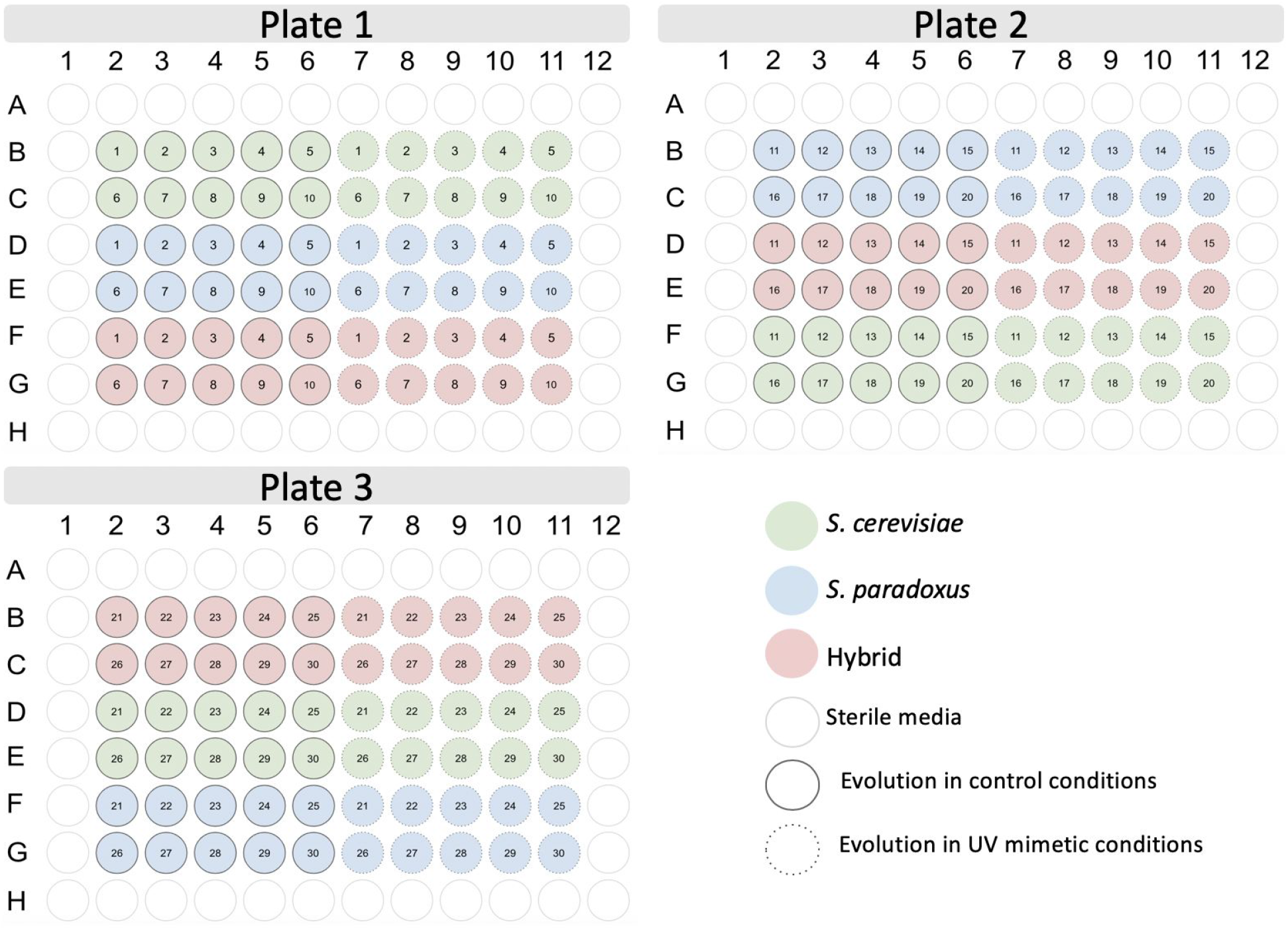
Design of the experiment. 30 parallel lines of each genotype were evolved under control and UV mimetic conditions in three different plates. The arrangement of the plates, with the genotypes changed position on each plate, allowed to limit the effect of position on the plate. The borders of the plates did not contain strains and were filled with sterile media to avoid border effects caused by evaporation.

**Figure S2.**
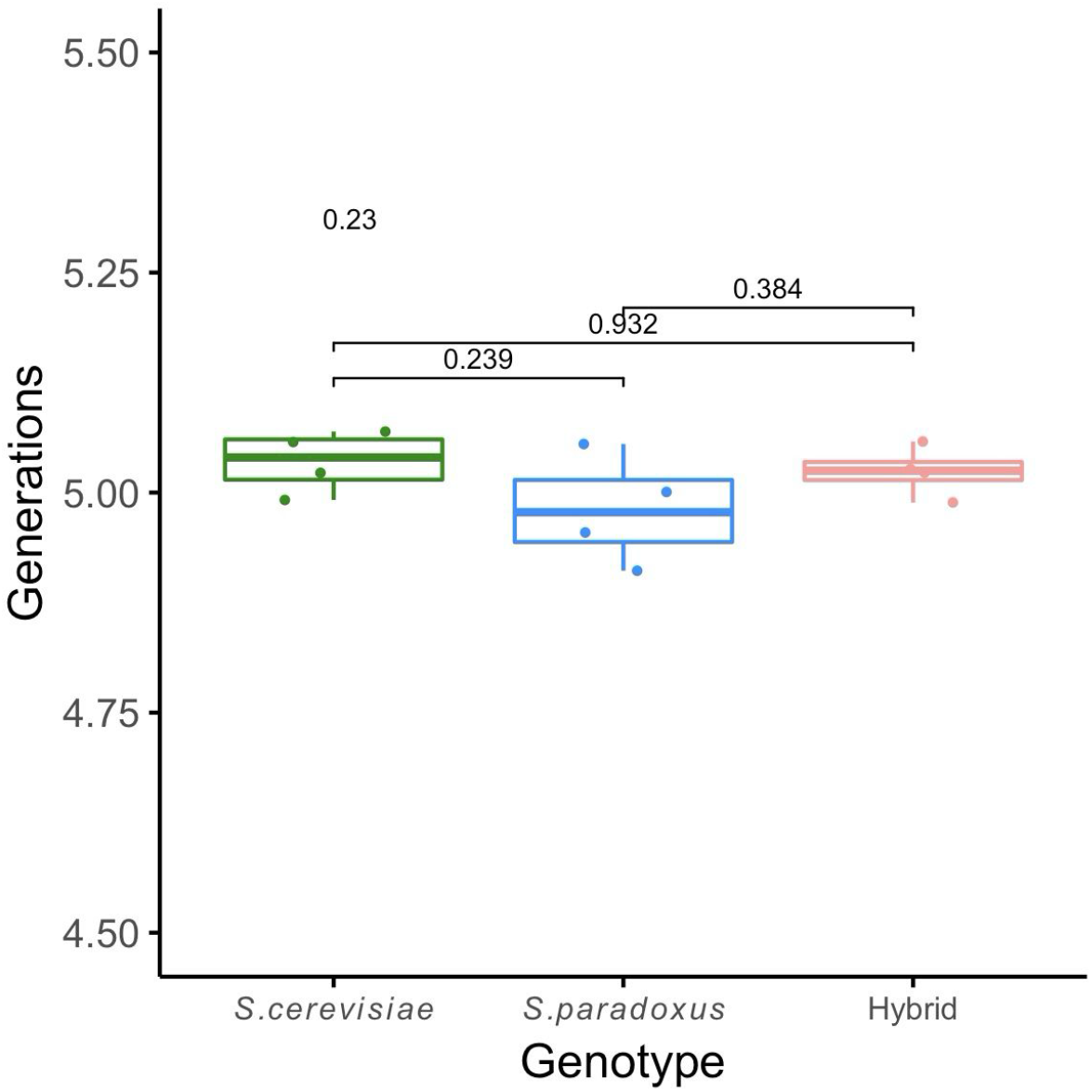
Number of generations. The number of generations was calculated with OD values at the beginning and the end of a cycle for the parental species and the hybrid. ANOVA and Tukey post-hoc pairwise comparisons were performed among genotypes (n = 4 populations for each genotype). P-value for ANOVA test (above) and Tukey post-hoc pairwise p-values are shown.

**Figure S3.**
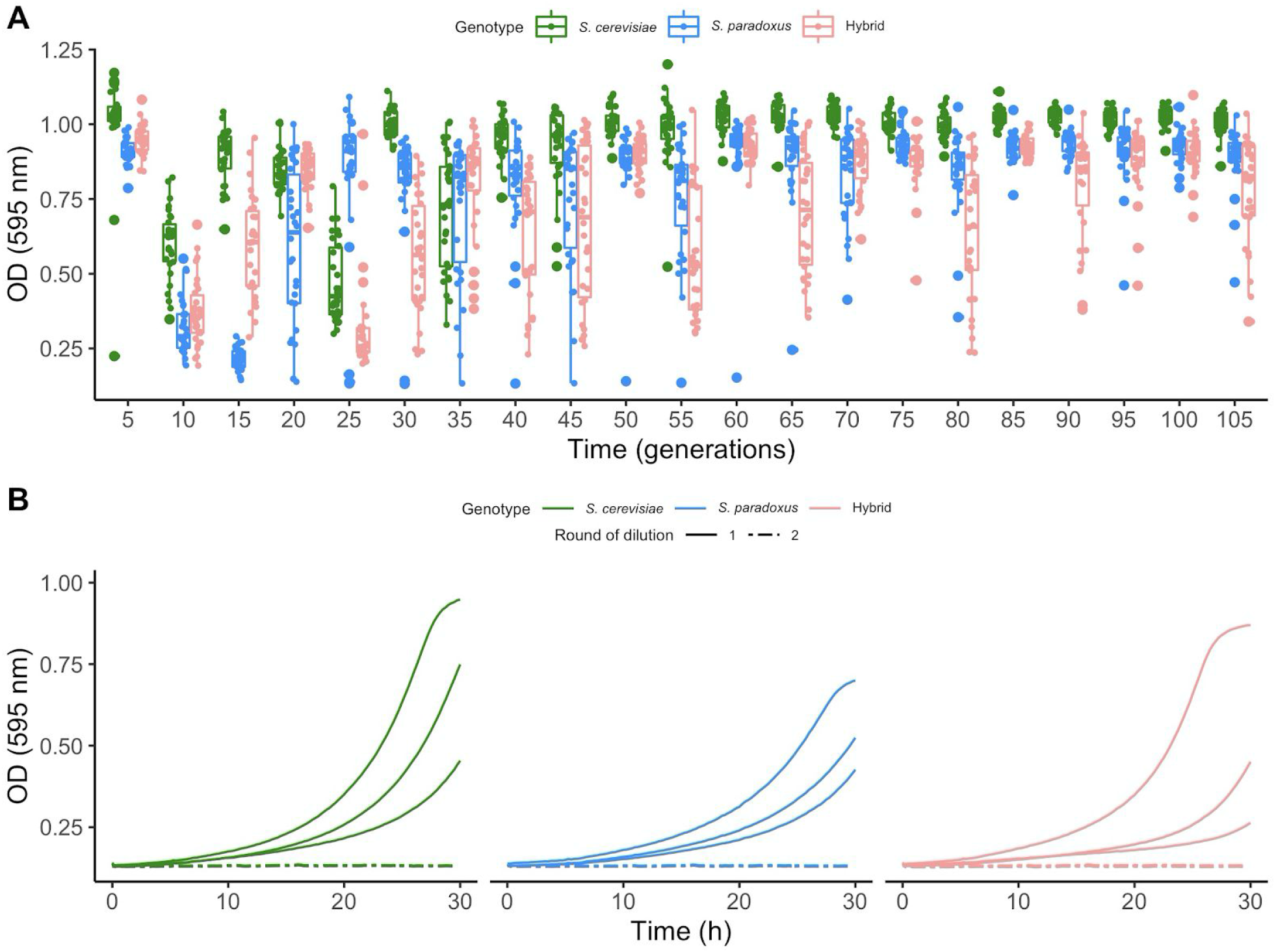
Cell sensitivity to the UV mimetic chemical increased during the two first cycles. A) Final optical density as a function of time (measured in generations) of each genotype in UV mimetic conditions (4 μM of 4-NQO) (n = 30 populations for each genotype). B) Optical density as a function of time (over 30 h) of each genotype in UV mimetic conditions (16 μM of 4-NQO) (n = 3 populations for each genotype).

**Figure S4.**
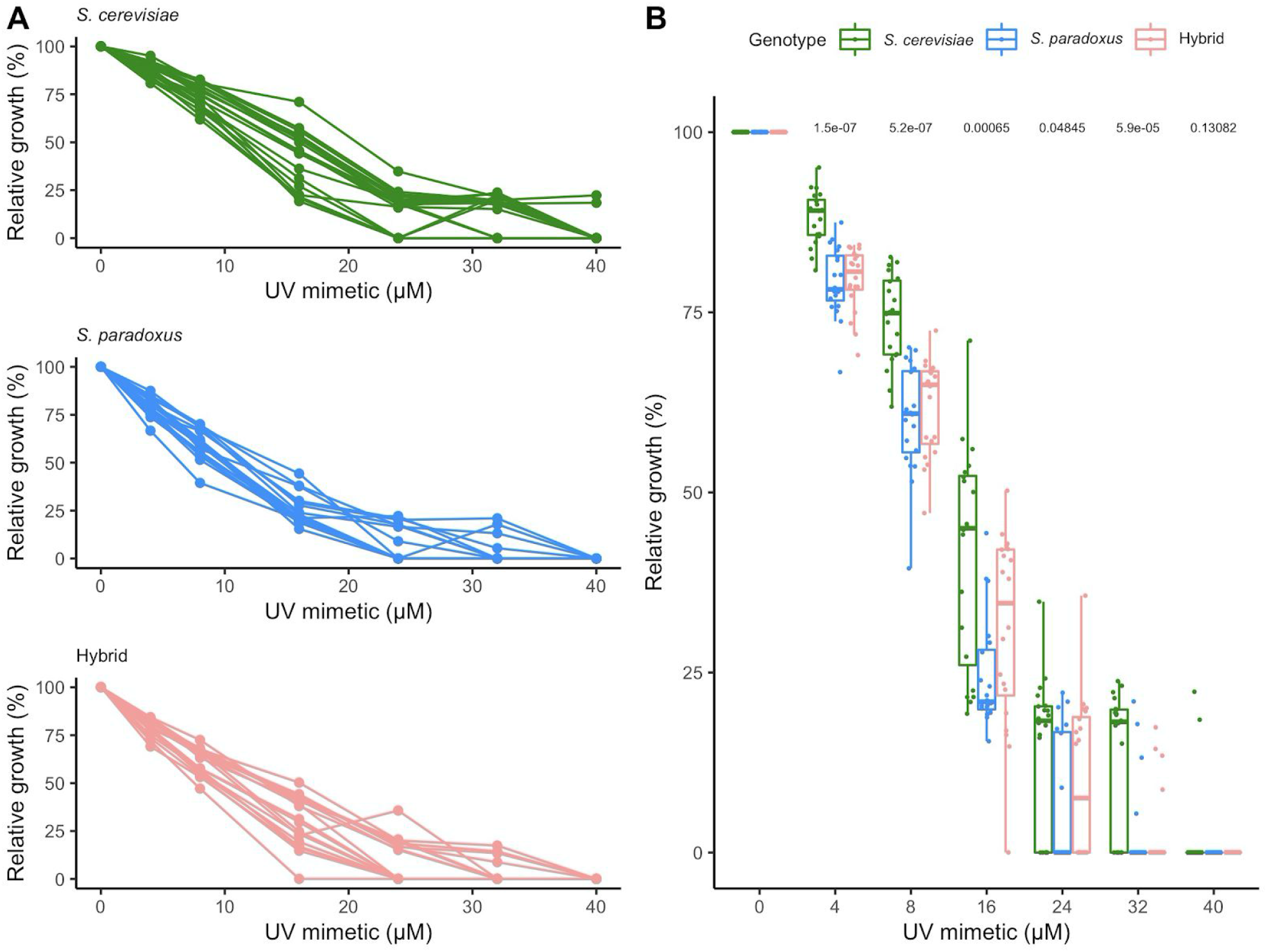
Effect of UV mimetic on growth. Relative growth is calculated by normalizing the growth rate in each concentration with respect to the growth rate of each strain in 0 μM of UV mimetic chemical (4-NQO) (n = 20 populations for each genotype). A) Growth of each population of each genotype in control (YPD) and in increasing UV mimetic concentrations (0 to 40 μM) B) Growth of each genotype in increasing UV mimetic concentrations (0 to 40 μM). ANOVAl of the relative growth (%) of the different genotypes as a function of the concentration was performed (n = 20 populations for each genotype). ANOVA test p-values for each concentration (above) are shown.

**Figure S5.**
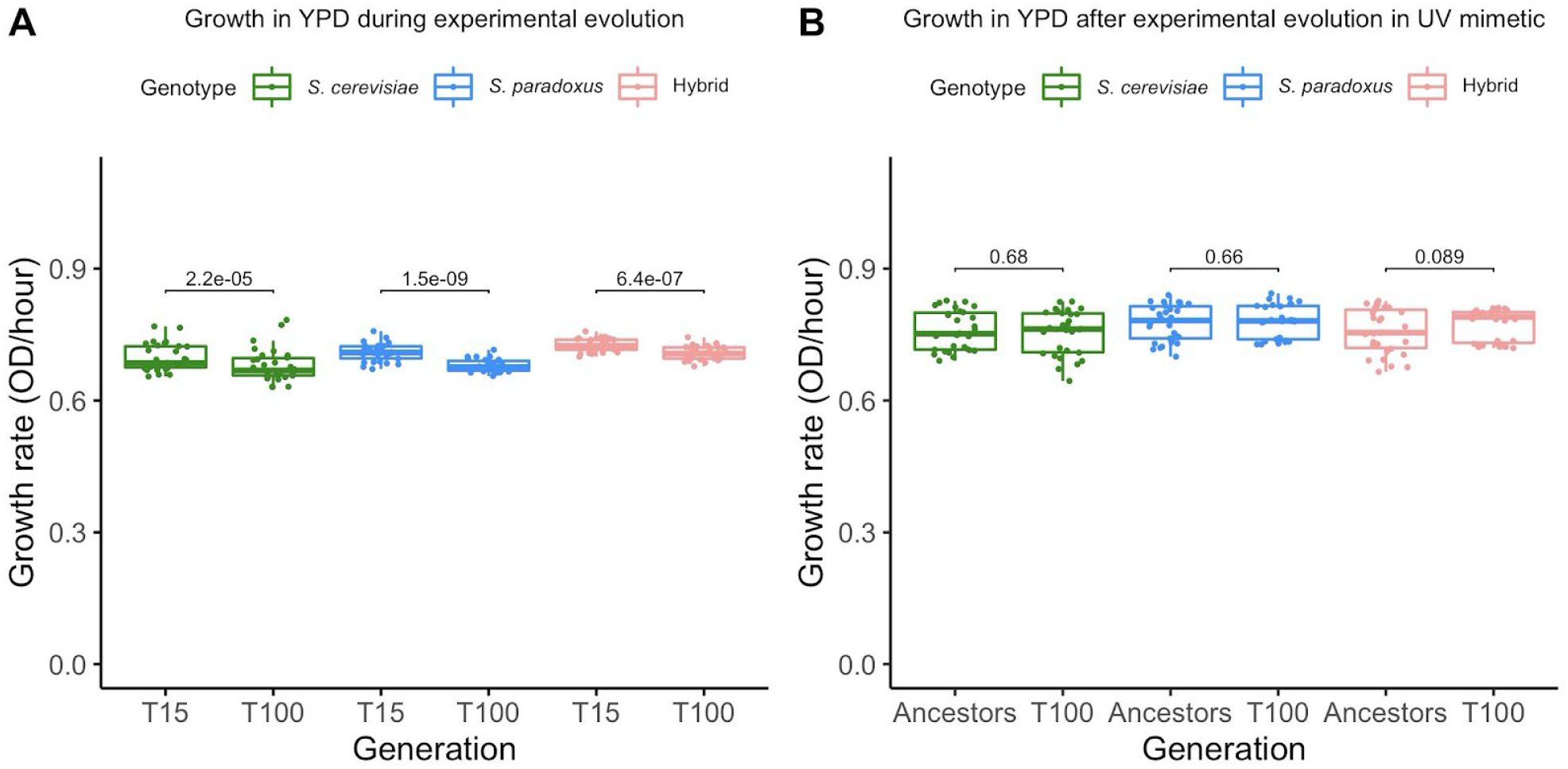
Growth of parents and hybrids in control conditions. A) Growth rates in control conditions (YPD) for the three genotypes evolving in UV mimetic at T15 and T100 (n = 30 populations for each genotype). Paired t-tests were performed between growth rate at 15 generations and at 100 generations for each genotype, pairing individual strains. P-values are shown (n = 30 populations for each genotype). B) Growth rates in control conditions for the three genotypes at ancestor state (from glycerol stock) and T100 (evolved in control from glycerol stock). Paired t-tests were performed between growth rate at ancestor state and at 100 generations for each genotype, pairing individual strains. P-values are shown (n = 30 populations for each genotype).

**Figure S6.**
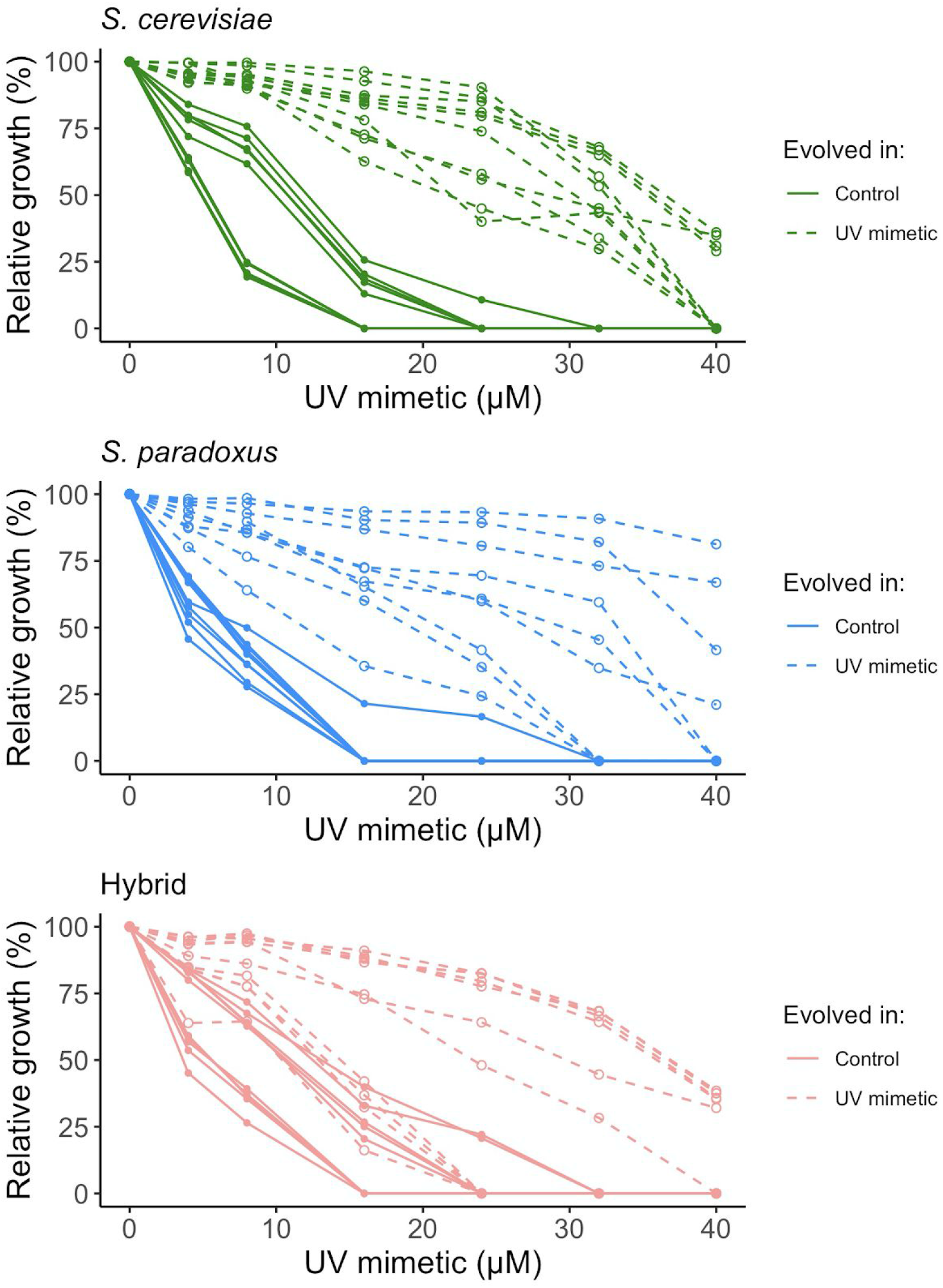
Adaptation to low concentration of UV mimetic leads to adaptation to higher concentrations. Growth of each population of each genotype evolved in UV mimetic or control conditions in increasing UV mimetic concentrations (0 to 40 μM) (n = 10 populations for each genotype).

**Figure S7.**
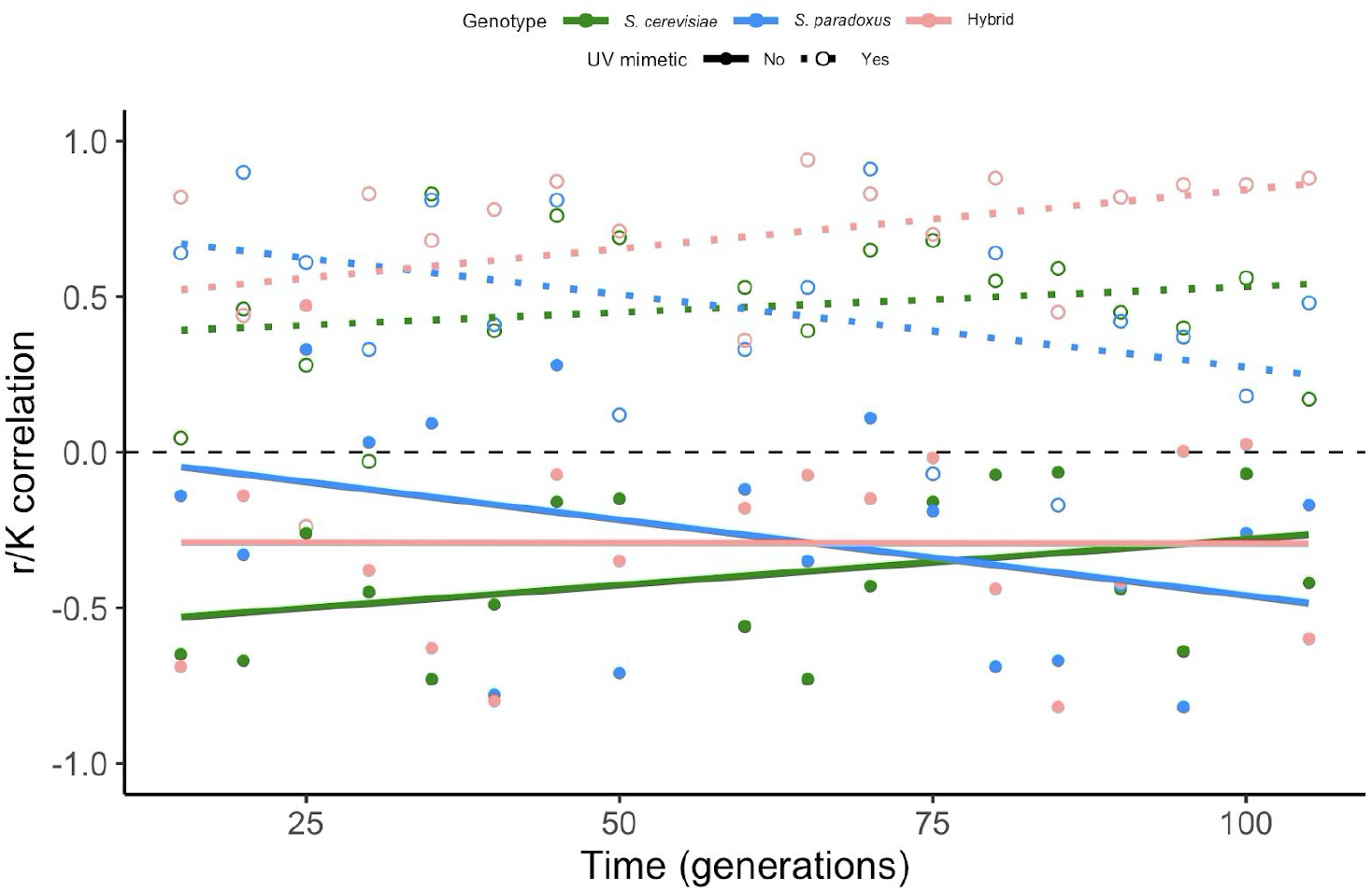
r/K correlations of parents and hybrids in control and UV mimetic conditions. Spearman’s rank correlations between growth rate (r) and carrying capacity (K) through generations for populations grown in control and in UV mimetic conditions. Each point represents the value of the Spearman’s rank correlation between r and K among the n = 30 populations for each genotype for a given day and a given condition.

**Figure S8.**
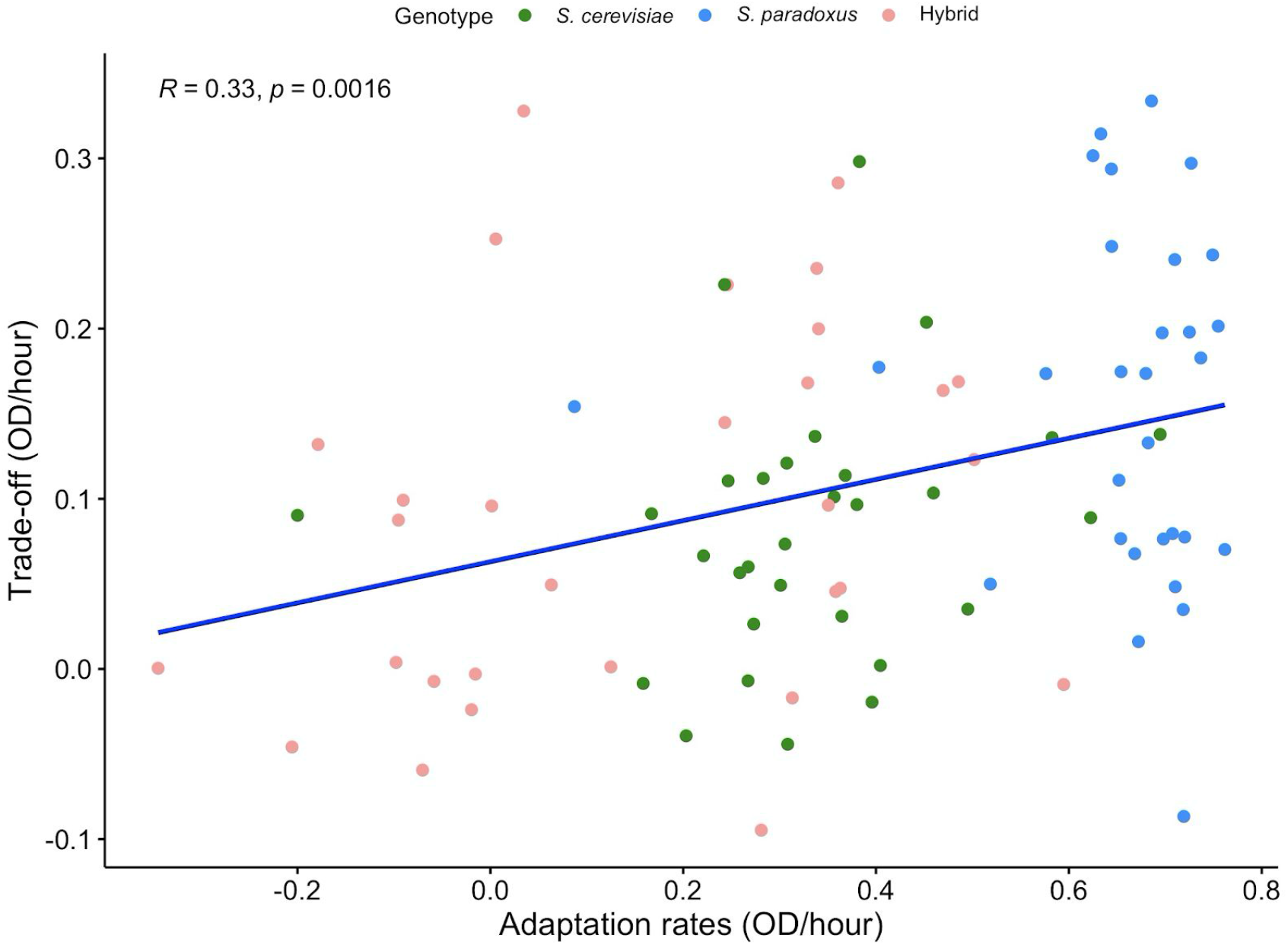
Trade-off depends on the extent of adaptation to UV mimetic conditions. Adaptation rates, expressed as fitness increase across the experiment, were obtained by subtracting the growth in UV mimetic conditions at T15 from the growth in UV mimetic conditions at T100. Trade-off estimates were obtained by subtracting the growth rate of the strains evolved in UV mimetic conditions (T100) grown in control conditions from the growth of the ancestors grown in control conditions. Spearman’s rank coefficient and associated p-value are shown.

**Table S1.** List of strains used in this study.

| Strain | Sampling site | Species | Mating type | Genotype | Reference Wild | Reference hoΔ |
| --- | --- | --- | --- | --- | --- | --- |
| MSH604 | Mont-St-Hilaire, QC | *S. paradoxus* | α | hoΔ::NAT | Leducq et al. 2014 | Leducq et al. 2016 |
| MSH604 | Mont-St-Hilaire, QC | *S. paradoxus* | a | hoΔ::HYG | Leducq et al. 2014 | Leducq et al. 2016 |
| LL13_054 | Rockport, MA | *S. cerevisiae* | α | hoΔ::NAT | Leducq et al. 2016 | Charron et al., 2019 |
| LL13_054 | Rockport, MA | *S. cerevisiae* | a | hoΔ::HYG | Leducq et al. 2016 | This study |

**Table S2.** Models fitted with their AIC score. Three models were used to fit the adaptation in UV mimetic conditions by expressing growth rate (rval) as a function of time and genotype.

| Model | AICc | Delta_AICc | ModelLik | AICcWt |
| --- | --- | --- | --- | --- |
| Asymptotic | -2745.955 | 0 | 1.00E+00 | 9.98E-01 |
| Logarithmic | -2733.785 | 12.17054 | 2.28E-03 | 2.27E-03 |
| Linear | -2656.771 | 89.1847 | 4.30E-20 | 2.27E-03 |

**Table S3.** Results of the linear model. “Scer” corresponds to *S. cerevisiae* and “Spar” corresponds to *S. paradoxus*.

| lm(formula = growth rate ~ generation * genotype) |  |  |  |
| --- | --- | --- | --- |
| Growth rate (OD/hour) |  |  |  |
|  | Estimate | Standard error | Pr(>\|t\|) |
| Intercept Generation + Genotype | 0.2418973 | 0.0108132 | < 2e-16 *** |
| Generation | 0.007298 | 0.0008129 | < 2e-16 *** |
| GenotypeScer | 0.0366239 | 0.0152922 | 0.0167 * |
| GenotypeSpar | 0.0044812 | 0.0153178 | 0.7699 |
| Generation:GenotypeScer | 0.008947 | 0.0011496 | 1.26e-14 *** |
| Generation:GenotypeSpar | 0.0061001 | 0.0011507 | 1.31e-07 *** |
| Degrees of freedom | 1612 |  |  |
| Adjusted R-squared | 0.4353 |  |  |

**Table S4.** Results of the Logarithmic model. The parameter “b” represents the increase in the growth rate while generations elapse for each genotype. “Scer” corresponds to *S. cerevisiae* and “Spar” corresponds to *S. paradoxus*.

| drm(rval ~ day, strain, fct = DRC.logCurve()) |  |  |  |
| --- | --- | --- | --- |
| Growth rate (OD/hour) |  |  |  |
|  | Estimate | Standard error | Pr(>\|t\|) |
| **b:Scer** | 0.1658994 | 0.0078283 | < 2.2e-16 *** |
| **b:Spar** | 0.1410741 | 0.0078422 | < 2.2e-16 *** |
| **b:Hybrid** | 0.075496 | 0.0078283 | < 2.2e-16 *** |
| **Comparison of parameter 'b'** |  |  |  |
| **Genotypes** | **p-value** |  |  |
| **Scer-Spar** | 0.0252 * |  |  |
| **Scer-Hybrid** | 6.516e-16 *** |  |  |
| **Spar-Hybrid** | 3.968e-09 *** |  |  |

**Table S5.** Results of the Asymptotic model. The parameter “c” is proportional to the increase in the growth rate while generations elapse for each genotype. The parameter “d” is the maximum attainable for each genotype. “Scer” corresponds to *S. cerevisiae* and “Spar” corresponds to *S. paradoxus*.

| drm(rval ~ day, strain, fct = AR.3()) |  |  |  |
| --- | --- | --- | --- |
| Growth rate (OD/hour) |  |  |  |
|  | Estimate | Standard error | Pr(>\|t\|) |
| **c:Scer** | 0.117474 | 0.033974 | 0.0005589 *** |
| **c:Spar** | -0.016485 | 0.071742 | 0.8182852 |
| **c:Hybrid** | 0.157355 | 0.037894 | 3.461e-05 *** |
| **d:Scer** | 0.625184 | 0.022568 | < 2.2e-16 *** |
| **d:Spar** | 0.489393 | 0.013855 | < 2.2e-16 *** |
| **d:Hybrid** | 0.391663 | 0.019385 | < 2.2e-16 *** |
| **Comparison of parameter 'c'** |  | **Comparison of parameter 'd'** |  |
| **Genotypes** | **p-value** | **Genotypes** | **p-value** |
| **Scer-Spar** | 0.09169 . | **Scer-Spar** | 3.286e-07 *** |
| **Spar-Hybrid** | 0.43338 | **Spar-Hybrid** | 7.558e-15 *** |
| **Spar-Hybrid** | 0.03229 * | **Spar-Hybrid** | 4.307e-05 *** |

## References

Adamo, G. M., Brocca, S., Passolunghi, S., Salvato, B., & Lotti, M. (2012). Laboratory evolution of copper tolerant yeast strains. Microbial Cell Factories, 11(1), 1.

Alix, K., Gérard, P. R., Schwarzacher, T., & Heslop-Harrison, J. S. P. (2017). Polyploidy and interspecific hybridization: partners for adaptation, speciation and evolution in plants. Annals of Botany, 120(2), 183–194.

Alves, M. J., Coelho, M. M., & Collares-Pereira, M. J. (2001). Evolution in action through hybridisation and polyploidy in an Iberian freshwater fish: a genetic review. Genetica, 111(1-3), 375–385.

Anderson, E. (1953). Introgressive Hybridization. Biological Reviews of the Cambridge Philosophical Society, 28(3), 280–307.

Anderson, Lee, Rushworth, Colautti, & Mitchell-Olds. (2013). Genetic trade-offs and conditional neutrality contribute to local adaptation. Molecular Ecology, 22(3), 699–708.

Baack, E. J., & Rieseberg, L. H. (2007). A genomic view of introgression and hybrid speciation. Current Opinion in Genetics & Development, 17(6), 513–518.

Ballerini, E. S. (2012). Natural hybridization as a catalyst of rapid evolutionary change. Rapidly Evolving Genes and Genetic Systems, 256–265.

Barbosa, R., Almeida, P., Safar, S. V. B., Santos, R. O., Morais, P. B., Nielly-Thibault, L., Leducq, J.-B., Landry, C. R., Gonçalves, P., Rosa, C. A., & Sampaio, J. P. (2016). Evidence of Natural Hybridization in Brazilian Wild Lineages of Saccharomyces cerevisiae. Genome Biology and Evolution, 8 (2), 317–329.

Becker, M., Gruenheit, N., Steel, M., Voelckel, C., Deusch, O., Heenan, P. B., McLenachan, P. A., Kardailsky, O., Leigh, J. W., & Lockhart, P. J. (2013). Hybridization may facilitate in situ survival of endemic species through periods of climate change. Nature Climate Change, 8(12), 1039–1043.

Bellon, J. R., Yang, F., Day, M. P., Inglis, D. L., & Chambers, P. J. (2015). Designing and creating Saccharomyces interspecific hybrids for improved, industry relevant, phenotypes. Applied Microbiology and Biotechnology, 99(20), 8597–8609.

Bleuven, C., & Landry, C. R. (2016). Molecular and cellular bases of adaptation to a changing environment in microorganisms. Proceedings. Biological Sciences / The Royal Society, 288(1841).

Burton, T. L., & Husband, B. C. (2000). Fitness differences among diploids, tetraploids, and their triploid progeny in Chamerion angustifolium: mechanisms of inviability and implications for polyploid evolution. Evolution; International Journal of Organic Evolution, 54(4), 1182–1191.

Caldwell, M. M., & Flint, S. D. (1994). Stratospheric ozone reduction, solar UV-B radiation and terrestrial ecosystems. Climatic Change, 28 (4), 375–394.

Chan, W. Y., Peplow, L. M., Menéndez, P., Hoffmann, A. A., & van Oppen, M. J. H. (2018). Interspecific Hybridization May Provide Novel Opportunities for Coral Reef Restoration. Frontiers in Marine Science, 5, 160.

Charron, G., Marsit, S., Hénault, M., Martin, H., & Landry, C. R. (2019). Spontaneous whole-genome duplication restores fertility in interspecific hybrids. Nature Communications, 10(1), 4126.

Cliften, P. F., Hillier, L. W., Fulton, L., Graves, T., Miner, T., Gish, W. R., Waterston, R. H., & Johnston, M. (2001). Surveying Saccharomyces genomes to identify functional elements by comparative DNA sequence analysis. Genome Research, 11(7), 1175–1186.

Cooper, V. S. (2014). The origins of specialization: insights from bacteria held 25 years in captivity. PLoS Biology, 12(2), e1001790.

Cooper, V. S., & Lenski, R. E. (2000). The population genetics of ecological specialization in evolving Escherichia coli populations. Nature, 407(6805), 736–739.

Couce, A., Caudwell, L. V., Feinauer, C., Hindré, T., Feugeas, J.-P., Weigt, M., Lenski, R. E., Schneider, D., & Tenaillon, O. (2017). Mutator genomes decay, despite sustained fitness gains, in a long-term experiment with bacteria. Proceedings of the National Academy of Sciences of the United States of America, 114(43) E9026–E9035.

de Visser, J. A. G. M., & Lenski, R. E. (2002). Long-term experimental evolution in Escherichia coli. XI. Rejection of non-transitive interactions as cause of declining rate of adaptation. BMC Evolutionary Biology, 2, 19.

Dhar, R., Sägesser R., Weikert, C., Yuan, J., & Wagner, A. (2011). Adaptation of Saccharomyces cerevisiae to saline stress through laboratory evolution. Journal of Evolutionary Biology, 24(5), 1135–1153.

Dion-Côté, A.-M., & Barbash, D. A. (2017). Beyond speciation genes: an overview of genome stability in evolution and speciation. Current Opinion in Genetics & Development, 47, 17–23.

Dobzhansky, T. & Pavlovsky, O. (1958). Interracial hybridization and breakdown of coadapted gene complexes in Drosophila paulistorum and Drosophila willistoni. Proceedings of the National Academy of Sciences, 44(6), 622–629.

Felkner, I. C., & Kadlubar F. (1968). Parallel between ultraviolet light and 4-nitroquinoline-1-oxide sensitivity in Bacillus subtilis. Journal of Bacteriology, 96(4), 1448–1449.

Gallego-Tévar, B., Curado, G., Grewell, B. J., Figueroa, M. E., & Castillo, J. M. (2018). Realized niche and spatial pattern of native and exotic halophyte hybrids. Oecologia, 188(3), 849–862.

Gallego-Tévar, B., Grewell, B. J., Futrell, C. J., Drenovsky, R. E., & Castillo, J. M. (2019). Interactive effects of salinity and inundation on native Spartina foliosa, invasive S. densiflora, and their hybrid from San Francisco Estuary, California. In Annals of Botany, 125(2), 377–389.

Gallego-Tévar, B., Rubio-Casal, A. E., de Cires, A., Figueroa, E., Grewell, B. J., & Castillo, J. M. (2018). Phenotypic plasticity of polyploid plant species promotes transgressive behaviour in their hybrids. AoB Plants, 10(5), ly055.

Gibson, B., & Liti, G. (2015). Saccharomyces pastorianus: genomic insights inspiring innovation for industry. Yeast, 32(1), 17–27.

Gilchrist, C., & Stelkens, R. (2019). Aneuploidy in yeast: Segregation error or adaptation mechanism? Yeast, 36(9), 525–539.

Gómez, J. M., González-Megías, A., Lorite, J., Abdelaziz, M., & Perfectti, F. (2015). The silent extinction: climate change and the potential hybridization-mediated extinction of endemic high-mountain plants. Biodiversity and Conservation, 24(8), 1843–1857.

Grant, B. R., & Grant, P. R. (1996). High Survival of Darwin's Finch Hybrids: Effects of Beak Morphology and Diets. Ecology, 77(2), 500–509.

Guerreiro, M. P. G. (2014). Interspecific hybridization as a genomic stressor inducing mobilization of transposable elements in Drosophila. Mobile Genetic Elements, 4(4), e34394.

Güldener, U., Heck, S., Fielder, T., Beinhauer, J., & Hegemann, J. H. (1996). A new efficient gene disruption cassette for repeated use in budding yeast. Nucleic Acids Research, 24(13), 2519–2524.

Hamilton, J. A., & Miller, J. M. (2016). Adaptive introgression as a resource for management and genetic conservation in a changing climate. Conservation Biology: The Journal of the Society for Conservation Biology, 30(1), 33–41.

Heil, C. S. S., Smukowski Heil, C. S., DeSevo, C. G., Pai, D. A., Tucker, C. M., Hoang, M. L., & Dunham, M. J. (2017). Loss of Heterozygosity Drives Adaptation in Hybrid Yeast. Molecular Biology and Evolution, 34(7), 1596–1612.

Herbst, R. H., Bar-Zvi, D., Reikhav, S., Soifer, I., Breker M., Jona, G., Shimoni, E., Schuldiner M., Levy, A. A., & Barkai, N. (2017). Heterosis as a consequence of regulatory incompatibility. BMC Biology, 15(1), 38.

Hessen, D. O. (1996). Competitive trade-off strategies in Arctic Daphnia linked to melanism and UV-B stress. Polar Biology, 16(8), 573–579.

Karpov, D. S., Spasskaya, D. S., Nadolinskaia, N. I., Tutyaeva, V. V., Lysov, Y. P., & Karpov, V. L. (2019). Deregulation of the 19S proteasome complex increases yeast resistance to 4-NQO and oxidative stress via upregulation of Rpn4- and proteasome-dependent stress responsive genes. FEMS Yeast Research, 19(2), foz002.

Khroustalyova, G., Giovannitti, G., Severini, D., Scherbaka, R., Turchetti, B., Buzzini, P., & Rapoport, A. (2019). Anhydrobiosis in yeasts: Psychrotolerant yeasts are highly resistant to dehydration. Yeast, 36(5), 375–379.

Landry, Hartl, & Ranz. (2007). Genome clashes in hybrids: insights from gene expression. Heredity, 99(5), 483–493.

Lawrence, C. W., & Christensen, R. (1976). UV mutagenesis in radiation-sensitive strains of yeast. Genetics, 82 (2), 207–232.

Leducq, J.-B., Nielly-Thibault, L., Charron, G., Eberlein, C., Verta, J.-P., Samani, P., Sylvester, K., Hittinger, C. T., Bell, G., & Landry, C. R. (2016). Speciation driven by hybridization and chromosomal plasticity in a wild yeast. Nature Microbiology, 1, 15003.

Lewontin, R. C., & Birch, L. C. (1966). Hybridization as a source of variation for adaptation to new environments. Evolution; International Journal of Organic Evolution, 20(3), 315–336.

Lexer, C., Welch, M. E., Durphy, J. L., & Rieseberg, L. H. (2003). Natural selection for salt tolerance quantitative trait loci (QTLs) in wild sunflower hybrids: implications for the origin of Helianthus paradoxus, a diploid hybrid species. Molecular Ecology, 12(5), 1225–1235.

Lidzbarsky, G. A., Shkolnik, T., & Nevo, E. (2009). Adaptive Response to DNA-Damaging Agents in Natural Saccharomyces cerevisiae Populations from “Evolution Canyon”, Mt. Carmel, Israel. PloS One, 4 (6), e5914.

Liu, J., Xu, X., & Deng, X. (2005). Intergeneric somatic hybridization and its application to crop genetic improvement. Plant Cell, Tissue and Organ Culture, 82 (1), 19–44.

Lopandic, K. (2018). Saccharomyces interspecies hybrids as model organisms for studying yeast adaptation to stressful environments. Yeast, 35(1), 21–38.

Lowe, W. H., Muhlfeld, C. C., & Allendorf, F. W. (2015). Spatial sorting promotes the spread of maladaptive hybridization. Trends in Ecology & Evolution, 30(8), 456–462.

Marcet-Houben, M., & Gabaldón, T. (2015). Beyond the Whole-Genome Duplication: Phylogenetic Evidence for an Ancient Interspecies Hybridization in the Baker's Yeast Lineage. PLoS Biology, 13(8), e1002220.

Marfil, C. F., Masuelli, R. W., Davison, J., & Comai, L. (2006). Genomic instability in Solanum tuberosum x Solanum kurtzianum interspecific hybrids. Genome / National Research Council Canada = Genome / Conseil National de Recherches Canada, 49(2), 104–113.

Marshall, J. E. A., Lakin, J., Troth, I., & Wallace-Johnson, S. M. (2020). UV-B radiation was the Devonian-Carboniferous boundary terrestrial extinction kill mechanism. Science Advances, 6 (22), eaba0768.

Martin, N. H., Bouck, A. C., & Arnold, M. L. (2006). Detecting adaptive trait introgression between Iris fulva and I. brevicaulis in highly selective field conditions. Genetics, 172(4), 2481–2489.

Mee, J. A., & Yeaman, S. (2019). Unpacking Conditional Neutrality: Genomic Signatures of Selection on Conditionally Beneficial and Conditionally Deleterious Mutations. The American Naturalist, 194(4), 529–540.

Mitchell, N., Owens, G. L., Hovick, S. M., Rieseberg, L. H., & Whitney, K. D. (2019). Hybridization speeds adaptive evolution in an eight-year field experiment. Scientific Reports, 9(1), 6746.

Morales, L., & Dujon, B. (2012). Evolutionary role of interspecies hybridization and genetic exchanges in yeasts. Microbiology and Molecular Biology Reviews: MMBR, 76(4), 721–739.

Murphy, H. A., Kuehne, H. A., Francis, C. A., & Sniegowski, P. D. (2006). Mate choice assays and mating propensity differences in natural yeast populations. Biology Letters, 2(4), 553–556.

Nolte, A. W., & Sheets, H. D. (2005). Shape based assignment tests suggest transgressive phenotypes in natural sculpin hybrids (Teleostei, Scorpaeniformes, Cottidae). Frontiers in Zoology, 2(1), 11.

Poelwijk, F. J., Kiviet, D. J., Weinreich, D. M., & Tans, S. J. (2007). Empirical fitness landscapes reveal accessible evolutionary paths. Nature, 445 (7126), 383–386.

Pulschen, A. A., Rodrigues, F., Duarte, R. T. D., Araujo, G. G., Santiago, I. F., Paulino-Lima, I. G., Rosa, C. A., Kato, M. J., Pellizari, V. H., & Galante, D. (2015). UV-resistant yeasts isolated from a high-altitude volcanic area on the Atacama Desert as eukaryotic models for astrobiology. MicrobiologyOpen, 4 (4), 574–588.

Replansky, T., Koufopanou, V., Greig, D., & Bell, G. (2008). Saccharomyces sensu stricto as a model system for evolution and ecology. Trends in Ecology & Evolution, 23(9), 494–501.

Rieseberg, L. H., Archer, M. A., & Wayne, R. K. (1999). Transgressive segregation, adaptation and speciation. Heredity, 83 (Pt 4), 363–372.

Rong-Mullins, X., Ayers, M. C., Summers, M., & Gallagher, J. E. G. (2018). Transcriptional Profiling of Saccharomyces cerevisiae Reveals the Impact of Variation of a Single Transcription Factor on Differential Gene Expression in 4NQO, Fermentable, and Nonfermentable Carbon Sources. G3, 8(2), 607–619.

Saitoh, K., Chen, W.-J., & Mayden, R. L. (2010). Extensive hybridization and tetrapolyploidy in spined loach fish. Molecular Phylogenetics and Evolution, 56(3), 1001–1010.

Salvadó, Z., Arroyo-López, F. N., Guillamón, J. M., Salazar, G., Querol, A., & Barrio, E. (2011). Temperature adaptation markedly determines evolution within the genus Saccharomyces. Applied and Environmental Microbiology, 77(7), 2292–2302.

Schierenbeck, K. A., & Ellstrand, N. C. (2009). Hybridization and the evolution of invasiveness in plants and other organisms. Biological Invasions, 11(5), 1093.

Schumer, M., Rosenthal, G. G., & Andolfatto, P. (2014). How common is homoploid hybrid speciation? Evolution; International Journal of Organic Evolution, 68(6), 1553–1560.

Schumer, M., Xu, C., Powell, D. L., Durvasula, A., Skov, L., Holland, C., Blazier J. C., Sankararaman, S., Andolfatto, P. Rosenthal, G. G., & Przeworski, M. (2018). Natural selection interacts with recombination to shape the evolution of hybrid genomes. Science, 360(6389), 656–660.

Seehausen, O. (2004). Hybridization and adaptive radiation. Trends in Ecology & Evolution, 19(4), 198–207.

Selmecki, A. M., Maruvka, Y. E., Richmond, P. A., Guillet, M., Shoresh, N., Sorenson, A. L., De, S., Kishony, R., Michor F., Dowell, R., & Pellman, D. (2015). Polyploidy can drive rapid adaptation in yeast. Nature, 519(7543), 349–352.

Siede, W., & Friedberg, E. C. (1990). Influence of DNA repair deficiencies on the UV sensitivity of yeast cells in different cell cycle stages. Mutation Research, 245(4), 287–292.

Soltis, P. S., & Soltis, D. E. (2009). The role of hybridization in plant speciation. Annual Review of Plant Biology, 60, 561–588.

Sprouffske, K., Aguilar-Rodríguez, J., Sniegowski, P. & Wagner, A. (2018). High mutation rates limit evolutionary adaptation in Escherichia coli. PLoS Genetics, 14(4), e1007324.

Sprouffske, K., & Wagner, A. (2016). Growthcurver: an R package for obtaining interpretable metrics from microbial growth curves. BMC Bioinformatics, 17, 172.

Stelkens, R. B., Brockhurst, M. A., Hurst, G. D. D., & Greig, D. (2014). Hybridization facilitates evolutionary rescue. Evolutionary Applications, 7(10), 1209–1217.

Sunshine, A. B., Payen, C., Ong, G. T.1 Liachko, I., Tan, K. M., & Dunham, M. J. (2015). The fitness consequences of aneuploidy are driven by condition-dependent gene effects. PLoS Biology, 13(5), e1002155.

Taddei, F., Radman, M., Maynard-Smith, J., Toupance, B., Gouyon, P. H., & Godelle, B. (1997). Role of mutator alleles in adaptive evolution. Nature, 387(6634), 700–702.

Vega, U., & Frey, K. J. (1980). Transgressive segregation in inter and intraspecific crosses of barley. Euphytica/ Netherlands Journal of Plant Breeding, 29(3), 585–594.

Waara, S., & Glimelius, K. (1995). The potential of somatic hybridization in crop breeding. Euphytica/Netherlands Journal of Plant Breeding, 85(1-3), 217–233.

Wahl, L. M., Gerrish, P. J., & Saika-Voivod, I. (2002). Evaluating the impact of population bottlenecks in experimental evolution. Genetics, 162(2), 961–971.

Wei, X., & Zhang, J. (2019). Environment-dependent pleiotropic effects of mutations on the maximum growth rate r and carrying capacity K of population growth. PLoS Biology, 17(1), e3000121.

Xie, Z., Wang, L., Wang, L., Wang, Z., Lu, Z., Tian, D., Yang, S., & Hurst, L. D. (2016). Mutation rate analysis via parent–progeny sequencing of the perennial peach. I. A low rate in woody perennials and a higher mutagenicity in hybrids. Proceedings of the Royal Society B: Biological Sciences, 283(1841), 20161016.

